# TDP-43 loss induces extensive cryptic polyadenylation in ALS/FTD

**DOI:** 10.1101/2024.01.22.576625

**Authors:** Sam Bryce-Smith, Anna-Leigh Brown, Puja R. Mehta, Francesca Mattedi, Alla Mikheenko, Simone Barattucci, Matteo Zanovello, Dario Dattilo, Matthew Yome, Sarah E. Hill, Yue A. Qi, Oscar G. Wilkins, Kai Sun, Eugeni Ryadnov, Yixuan Wan, NYGC ALS Consortium, Jose Norberto S. Vargas, Nicol Birsa, Towfique Raj, Jack Humphrey, Matthew Keuss, Michael Ward, Maria Secrier, Pietro Fratta

## Abstract

Nuclear depletion and cytoplasmic aggregation of the RNA-binding protein TDP-43 is the hallmark of ALS, occurring in over 97% of cases. A key consequence of TDP-43 nuclear loss is the de-repression of cryptic exons. Whilst TDP-43 regulated cryptic splicing is increasingly well catalogued, cryptic alternative polyadenylation (APA) events, which define the 3’ end of last exons, have been largely overlooked, especially when not associated with novel upstream splice junctions. We developed a novel bioinformatic approach to reliably identify distinct APA event types: alternative last exons (ALE), 3’UTR extensions (3’Ext) and intronic polyadenylation (IPA) events. We identified novel neuronal cryptic APA sites induced by TDP-43 loss of function by systematically applying our pipeline to a compendium of publicly available and in house datasets. We find that TDP-43 binding sites and target motifs are enriched at these cryptic events and that TDP-43 can have both repressive and enhancing action on APA. Importantly, all categories of cryptic APA can also be identified in ALS and FTD post mortem brain regions with TDP-43 proteinopathy underlining their potential disease relevance. RNA-seq and Ribo-seq analyses indicate that distinct cryptic APA categories have different downstream effects on transcript and translation. Intriguingly, cryptic 3’Exts occur in multiple transcription factors, such as *ELK1*, *SIX3,* and *TLX1,* and lead to an increase in wild-type protein levels and function. Finally, we show that an increase in RNA stability leading to a higher cytoplasmic localisation underlies these observations. In summary, we demonstrate that TDP-43 nuclear depletion induces a novel category of cryptic RNA processing events and we expand the palette of TDP-43 loss consequences by showing this can also lead to an increase in normal protein translation.

## Introduction

Cytoplasmic aggregates and nuclear depletion of TDP-43 are pathological hallmarks of a spectrum of neurodegenerative diseases, including over 97% of amyotrophic lateral sclerosis (ALS) cases^1^, 45% of frontotemporal dementia (FTD)^2^ and over 50% of Alzheimer’s disease cases^3^. Under normal conditions, TDP-43 is a predominantly nuclear protein with multiple roles in regulation of RNA processing and metabolism, including alternative splicing, alternative polyadenylation (APA)^4–6^, and transport^7^. Significant attention has been drawn to TDP-43’s ability to repress the inclusion of pre-mRNA sequences in mature transcripts^8^: loss of nuclear TDP-43 leads to the inclusion of ‘cryptic’ exons in mature transcripts both *in vitro* and in post mortem tissue^7^, contributing to disease progression^10,11^. Cryptic exons can lead to protein loss through RNA degradation by nonsense mediated decay^12^, or can be translated to produce cryptic peptides^13,14^.

Cleavage and polyadenylation defines the 3′end of last exons and subsequently mature transcripts^15^. Up to 70% of human protein-coding and long non-coding RNA genes can undergo polyadenylation at multiple locations in the gene body (alternative polyadenylation, APA), and can be subdivided in three main category of events: alternative last exons (ALE), 3’UTR extensions (3’Ext) and intronic polyadenylation events (IPA). In alternative last exons (ALE), the polyA usage is determined by an upstream alternative splice junction which defines an alternative last exon. In 3’Ext events, APA sites are independent of splice junctions and occur within 3’UTR regions and affect 3’UTR sequence and length, which is implicated in the regulation of transcript stability, localisation and translation^16^. Finally in IPA events, APA occurs within introns giving rise to transcripts with different protein coding potential and can affect full-length protein dosage^17,18^.

TDP-43 regulated cryptic APA has not been systematically explored in a neuronal context. Here, we report widespread cryptic APA upon TDP-43 depletion in cell models, including events which were not previously detected with conventional splicing analyses. A substantial number are expressed in post-mortem ALS & ALS/FTD tissue with TDP-43 loss, underlining their potential involvement in pathogenic mechanisms and/or utility as biomarkers of TDP-43 pathology. We focus on a novel class of 3’Ext APA and show they can lead to increased translation levels. Moreover, we use metabolic labelling to demonstrate that such cryptic 3’Ext are associated with increased RNA stability, and in the case of *ELK1*, coincide with increased cytoplasmic RNA localisation. Our data therefore identifies a novel consequence for cryptic RNA processing, and shows that in addition to leading to protein reduction or the formation of altered proteins, this can also lead to overexpression of normal proteins, and an increase in their function.

## Results

### Identification of cryptic alternative polyadenylation events induced by TDP-43 loss

While TDP-43’s role in regulating APA and cryptic splicing is well-known, cryptic APA occurring upon TPD-43 loss-of-function has yet to be explored. In order to comprehensively address this question, we curated a compendium of publicly available and newly generated bulk RNA-seq datasets with TDP-43 depletion (**Supplementary Table 1**). We assembled a computational pipeline to identify novel last exons from RNA-seq data, which defines last exon frames using StringTie^19^, and then filters and categorises as spurious predicted 3’ends lacking the presence of reference polyA sites^20^ or a conserved polyA signal hexamer^21^ (**Fig. 1A**). Isoform level quantification was performed using Salmon^22^, and differential usage between experimental conditions was assessed using DEXSeq^23^.

**Figure 1-.**
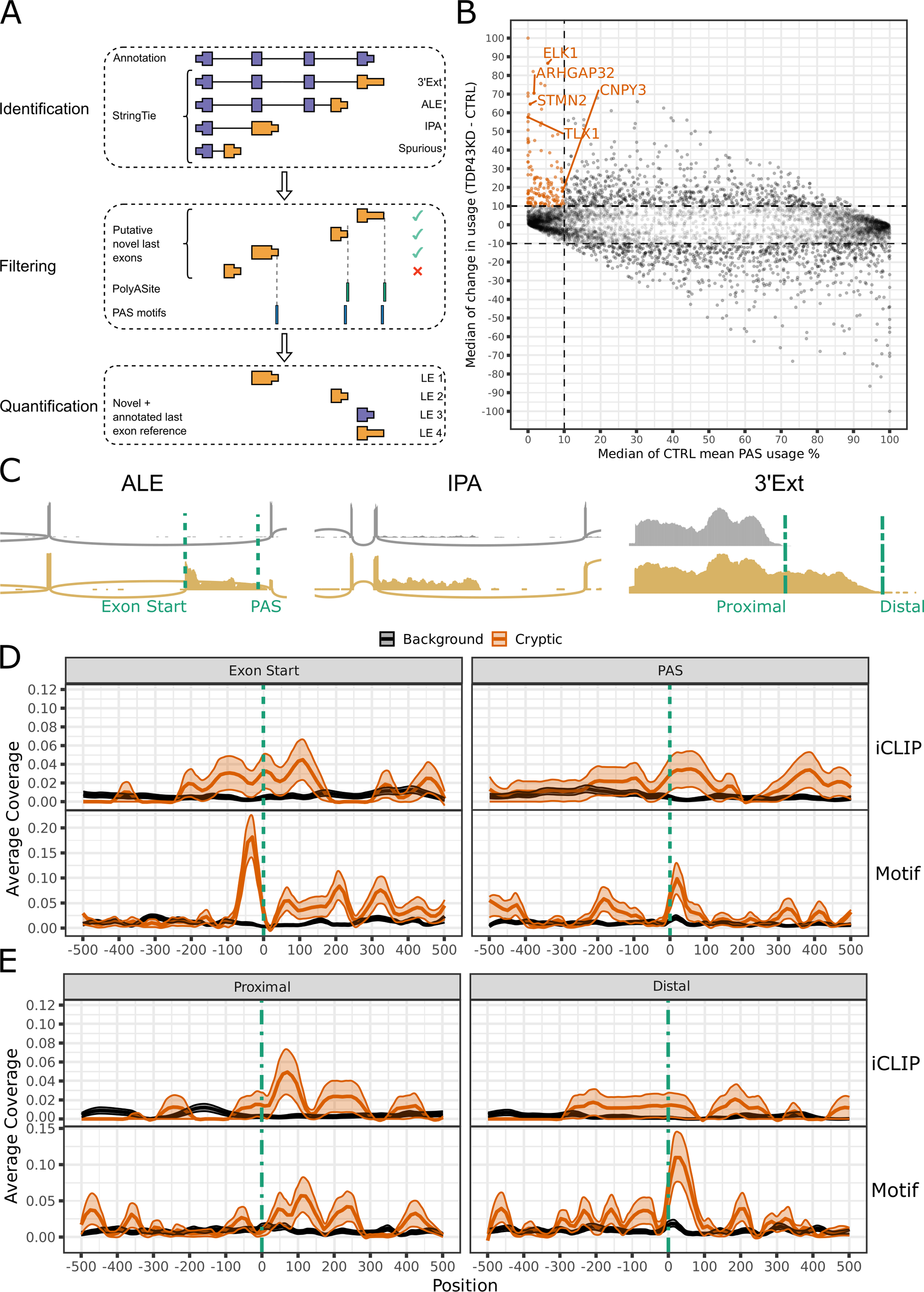
TDP-43 depletion induces cryptic APA in a compendium of in vitro TDP-43 datasets. A) Schematic demonstration of the computational pipeline to detect, quantify and infer differential usage of last exons from bulk RNA-seq data. De-novo transcripts are assembled using StringTie^19^ and subsequently filtered for a mean TPM > 1 within each experimental condition. Last exons are extracted from transcript models and compared to reference annotation (purple) to identify putative novel last exons (orange). Putative last exons are filtered for predicted 3’ends < 100nt from sites in the PolyASite^20^ database, or rescued if a conserved polyA signal hexamer^21^ can be identified within the last 100nt of the last exon. Novel and annotated last exons were subsequently quantified using Salmon^22^ and assessed for differential usage using DEXSeq^23^. For further details see Methods. B) Last exons responsive to TDP-43 depletion. All points represent a last exon passing a Benjamini-Hochberg adjusted p-value < 0.05 threshold in at least one dataset. Where a last exon passes the threshold in multiple datasets, the median values across datasets are calculated to represent the basal usage and change in usage upon TDP-43 depletion. Last exons passing our cryptic threshold are highlighted in orange (Benjamini-Hochberg adjusted p-value < 0.05, mean usage in control cells < 10 % and change in usage between TDP-43 knockdown and control (‘TDP43KD’ - ‘CTRL’) > 10 %). C) Example RNA-seq coverage traces control (grey) and TDP-43 knockdown (gold) i3Neuron samples for cryptic ALE (*ARHGAP32*), IPA (*ANKRD27*) and 3’Ext (*TLX1*) events. Dashed lines indicate landmarks around which TDP-43 binding is assessed in (C) and (D). *ARHGAP32* and *ANKRD27* are encoded on the reverse strand but flipped to read 5’-3’ for visualisation purposes. D) TDP-43 binding maps around boundaries of ALEs. (Top) TDP-43 iCLIP RNA maps around the first nucleotide of the last exon (‘Exon Start’) and the polyA site (‘PAS’) of ALEs. The solid lines represent the mean coverage (equivalent to fraction coverage) of relative positions upstream (negative values) and downstream (positive values) of the landmark from pooled TDP-43 iCLIP peaks from SH-SY5Y cells (n=2) in background (black, n=929) and cryptic ALEs (orange, n=92). Shaded intervals represent +/− 1 SE for the corresponding coloured group. (Bottom) TDP-43 motif maps around the first nucleotide of the last exon (‘Exon Start’) and the polyA site (‘PAS’) of ALEs. The solid lines represent the mean coverage by YG-containing hexamers (Supplementary Fig. 3A) of relative positions upstream (negative values) and downstream (positive values) of the landmark in background (black, n=929) and cryptic ALEs (orange, n=92). E) TDP-43 binding maps around alternative polyA sites of 3’Exts. (Top) TDP-43 iCLIP RNA maps around the proximal (‘Proximal’) and distal (‘Distal’) polyA site of 3’Ext events. As in D), but background (black, n=798) and cryptic regions (orange, n=86) are obtained for 3’Ext events. (Bottom) TDP-43 motif maps around the proximal (‘Proximal’) and distal (‘Distal’) polyA site of 3’Ext events. As in D), but background (black, n=798) and cryptic regions (orange, n=86) are obtained for 3’Ext events.

We subdivided our events into three main categories: ALEs, IPAs and 3’Ext (**Fig. 1A**). APA events were widespread and we defined cryptic APA events as ones with <10% mean usage in controls and >10% usage change after TDP-43 knockdown. We identified 227 cryptic APAs to be present in at least 1 dataset (adjusted P < 0.05, **Fig. 1B, Supplementary Fig. 1, Supplementary Table 2**). Cryptic ALEs (n=92) included previously identified cryptic exons such as *STMN2*, *ARHGAP32,* and *RSF1* (**Fig. 1B**). 108 3’UTR cryptics were identified, of which 86 are novel 3’UTR extensions (3’Ext; e.g. *TLX1,* **Fig. 1C**) and 20 were 3’UTR shortening events (proximal 3’Ext). 20 IPA events were also detected, including *CNPY3* which was identified and experimentally validated in an orthogonal bioinformatics approach (see Arnold et al., co-submitted). The remaining 9 events could not be uniquely assigned to ALEs or IPAs based on annotation, and are defined as ‘complex’.

70% (159/227) of cryptic APAs were detected as significant in a single dataset (**Supplementary Fig. 2A**), but we found that the majority (138) satisfied cryptic expression criteria (<10% mean usage in controls and >10% usage change after TDP-43 knockdown) across datasets. 51 APAs were consistently below 10% usage threshold in controls, but did not sufficiently increase following TDP-43 depletion to meet the cryptic criteria definition across datasets. 28 APAs showed instead a significant increase upon TDP-43 loss across datasets, but had >10% median usage in controls, therefore placing them outside the cryptic criteria, but demonstrating consistent regulation by TDP-43 (**Supplementary Fig. 2B**). Altogether, this data highlights a widespread presence of cryptic APA upon TDP-43 loss.

### TDP-43 binding can both repress and enhance polyA site selection

Next, we investigated TDP-43 binding patterns around cryptic APAs using TDP-43 iCLIP data generated in SH-SY5Y cells^10^. We focussed on ALEs and 3’Ext events as the low number of IPA and proximal 3’Ext events (n=20 in both cases) did not allow reliable binding profile inferences. TDP-43 binding was enriched around the splice acceptor of cryptic ALEs as previously described in cryptic splice junctions and downstream of the cryptic polyadenylation site (PAS) of ALEs **(Fig. 1D)**, supporting TDP-43 acting as a repressor of both splicing and polyadenylation. Intriguingly, TDP-43 binding was also enriched immediately downstream of the annotated proximal PAS of 3’Ext events (**Fig. 1E**), supporting a role for TDP-43 in enhancing polyA usage consistent with previous reports of TDP-43 binding with respect to regulated PAS^5^.

iCLIP data, typically generated in control cells, is not sensitive in detecting binding to cryptic 3’Ext regions, as these events can be only detected at very low levels with physiological TDP-43 presence. We therefore sought to corroborate our findings by adapting PEKA^24^ to infer de-novo hexamer enrichment relative to cryptic landmarks. Previously defined hexamers enriched around TDP-43 iCLIP binding sites^6^ (**Supplementary Fig. 3A**) were overrepresented among the most enriched hexamers proximal to all cryptic landmarks, with the strongest signal overall observed at both the 3’ss and PAS of ALE events (**Supplementary Fig. 3B**). To assess the concordance with iCLIP binding profiles, we visualised the positional coverage of the hexamer group most strongly associated with TDP-43 binding^6^. We observed a striking peak immediately upstream of ALE splice acceptors, consistent with the previously observed mechanism of *STMN2* cryptic exon repression^25^ (**Fig. 1D**). Enriched signal was also observed immediately downstream of the distal PAS of 3’Exts (**Fig. 1E)** and the PAS of ALEs (**Fig. 1D**). Overall, our data support a direct role for TDP-43 binding in both enhancing and repressing PAS usage, therefore leading to cryptic APA upon TDP-43 loss.

### TDP-43 cryptic APA is detectable in post-mortem ALS/FTD tissues

We next investigated whether the cryptic APA detected *in vitro* occurred also in post-mortem central nervous system (CNS) tissue samples affected by TDP-43 proteinopathy. We initially focused on neuronal nuclei sorted into TDP-43 positive and TDP-43 negative populations^26^.

60 cryptic APA events were more highly expressed in TDP-43 depleted nuclei. All APA event types were represented in this list (**Fig. 2A**), with ALEs (28) and 3’Exts (27) representing the majority of enriched events. Our analysis confirms previously reported cryptic ALEs with patient specificity such as in *STMN2*^27^. A number of 3’Ext also show enrichment in TDP-43 negative nuclei in a similar magnitude to *STMN2* (median increased usage of 69 %), most notably *ELK1 (76 %)* and *RBM27 (57 %)* (**Fig. 2A**). Five IPA events meet our enrichment criteria (**Fig. 2A**), including *USP31*, which was identified in a targeted assay of sporadic ALS motor cortex tissue^28^. However, IPA events were generally more weakly enriched in TDP-43 depleted nuclei compared to 3’Ext and ALE events. Altogether, this analysis shows that cryptic APA is detectable in post-mortem ALS/FTD CNS.

**Figure 2-.**
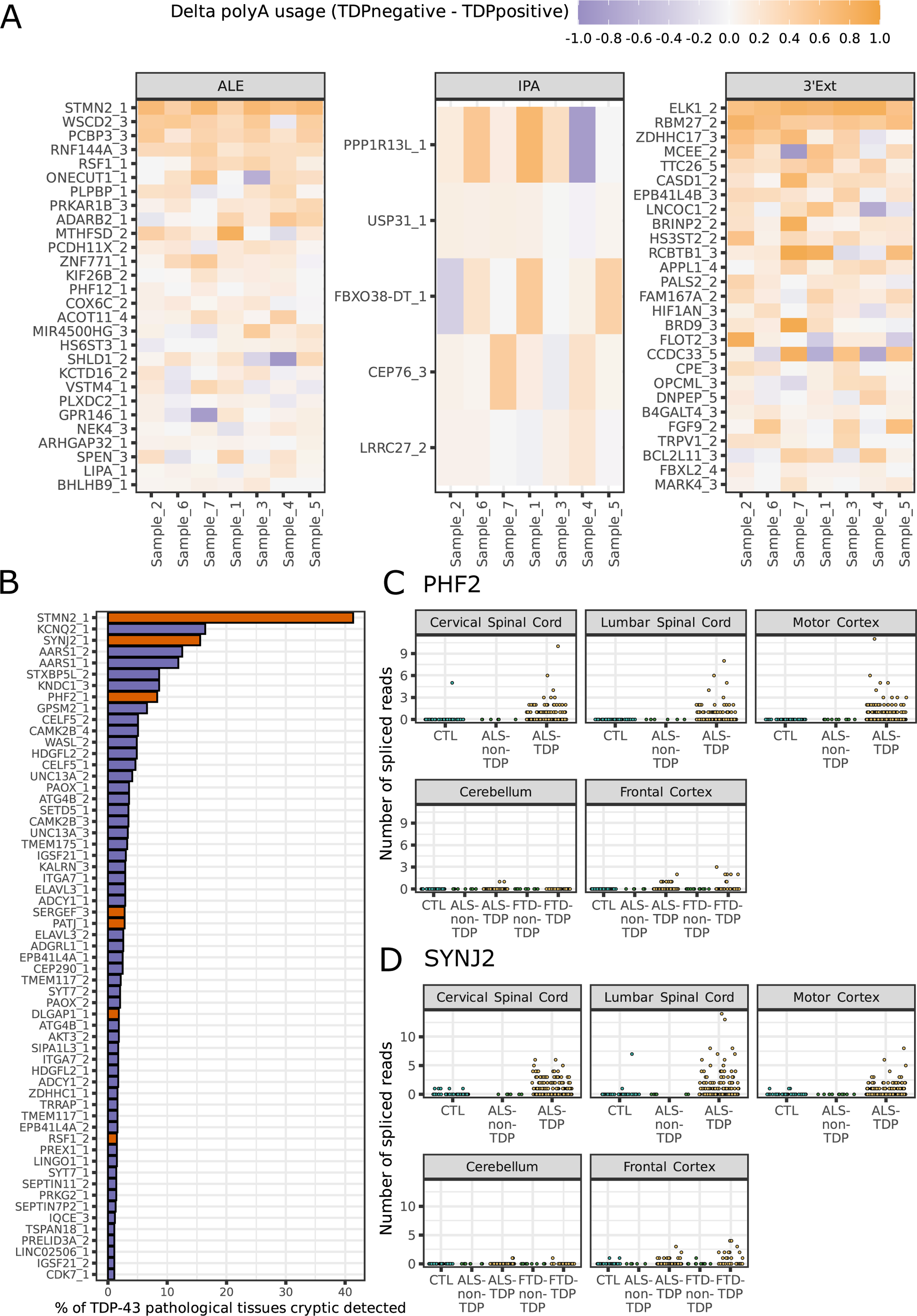
Cryptic last exons are detected in post-mortem ALS-FTD RNA-seq datasets. A) Heatmap of cryptic polyadenylation site usage in post-mortem FACS-seq data^26^. Cells are coloured according to the magnitude of sample-wise difference in usage between TDP-43 depleted (TDPnegative) and TDP-43 positive (TDPpositive) cells. Rows represent individual cryptic last exons from in-vitro that passed enrichment criteria (median sample-wise difference in usage (TDPnegative - TDPpositive) > 5 %) and are arranged in descending order of the difference in usage within each event type. Columns represent individual patients within the cohort. B) Selectively expressed cryptic ALEs (orange) and splicing events^13^ (purple) in tissues and samples with TDP-43 proteinopathy in the New York Genome Centre (NYGC) ALS Consortium dataset. Events are considered detected if at least 2 junction reads were detected in a sample. C) Detection of spliced reads for the cryptic ALE in *PHF2* across samples in the NYGC ALS Consortium dataset. ‘CTL’ denotes control samples. Colour indicates whether disease subtype and region is expected (orange) or not expected (green) to have TDP-43 pathology and cryptic spliced read expression. D) As in C), but for cryptic ALE in *SYNJ2*.

Next, we used the New York Genome Centre (NYGC) ALS consortium RNA-seq dataset to assess cryptic APA in a larger cohort of CNS cases with or without TDP-43 pathology. Cryptic 3’Exts often demonstrated low basal expression in control samples in our *in vitro* datasets, confounding the detection in post-mortem bulk RNA-seq datasets, where only a very small proportion of cells is expected to have TDP-43 pathology. IPA detection is further complicated by the fact that normal pre-mRNA reads also map to IPA regions creating significant noise in bulk RNA-seq. We therefore focussed on ALEs, where detection of the associated upstream cryptic splice junctions provide direct evidence of expression. As cryptic ALEs are expected to be dependent on nuclear TDP-43 depletion, we defined criteria based on spliced read detection to identify cryptic events with specific expression in tissues and disease subtypes where TDP-43 pathology is present. 7/118 cryptic ALE junctions fulfilled specificity criteria (**Supplementary Table 3**), in contrast to 56/313 cryptic splicing events collated from i3Neurons with TDP-43 knockdown^13^ (**Fig. 2B**). *STMN2* was most frequently detected in tissues with expected TDP-43 proteinopathy, and several other ALEs were amongst the most frequently detected specific cryptic events, including *SYNJ2* (3rd, **Fig. 2C**) and *PHF2* (8th, **Fig. 2D**). Altogether, this suggests that cryptic APAs are detectable in post-mortem tissue affected by TDP-43 pathology, highlighting their potential relevance in loss-of-function disease mechanisms and their promising utility as biomarkers.

### Cryptic APA events have variable effects on differential expression

Cryptic splicing events impact expression, often leading to a reduction in transcript levels^9–11^. We therefore assessed the effect of cryptic APAs on their own transcripts in i3Neurons^13^ (**Supplementary Fig. 4A**), and found that the majority of events (86/126) coincide with a significant change in expression, equally split between significant upregulation and downregulation. When subdivided further into cryptic APA categories, no category showed a clear bias for upregulation or downregulation (19/34 3’Ext, 17/37 ALE and 6/10 IPA genes are downregulated). This suggests that cryptic APAs are associated with differential expression, but have variable effects on transcript levels.

### Cryptic 3’UTR extensions in transcription factor RNAs lead to increased translation and function

Regulation of both ALE and 3’Ext usage has been demonstrated to impact protein abundance through distinct mechanisms^29,30^, but differential RNA abundance does not necessarily imply a coordinated change in protein levels. To assess whether changes in gene expression were also reflected in translation levels, we performed differential translation analysis of Ribo-seq data generated from i3Neurons with TDP-43 depletion^13^.

Only a minority of cryptic APA-containing genes (26/126) showed significant changes in overall translation levels (**Supplementary Table 4**), of which 24 are concordantly altered in both Ribo-seq and RNA-seq abundance upon TDP-43 KD (**Fig. 3A,3B**). Notably, the differentially translated subset appeared to stratify by APA category: whilst ALEs are downregulated, all four significant 3’Exts, which also showed increased RNA abundance (**Fig. 3A**), had significantly increased translation (**Fig. 3B**). Gene set enrichment analysis (GSEA)^31,32^ confirmed that cryptic ALE and 3’Ext genes are significantly associated with decreased (normalised enrichment score (NES) −2.09, padj 2.31e-6) and increased translation (NES 1.54, padj 0.03) respectively, whilst IPA genes show no significant association in either direction (NES −1.09, padj 0.36, **Supplementary Fig. 4B**).

**Figure 3-.**
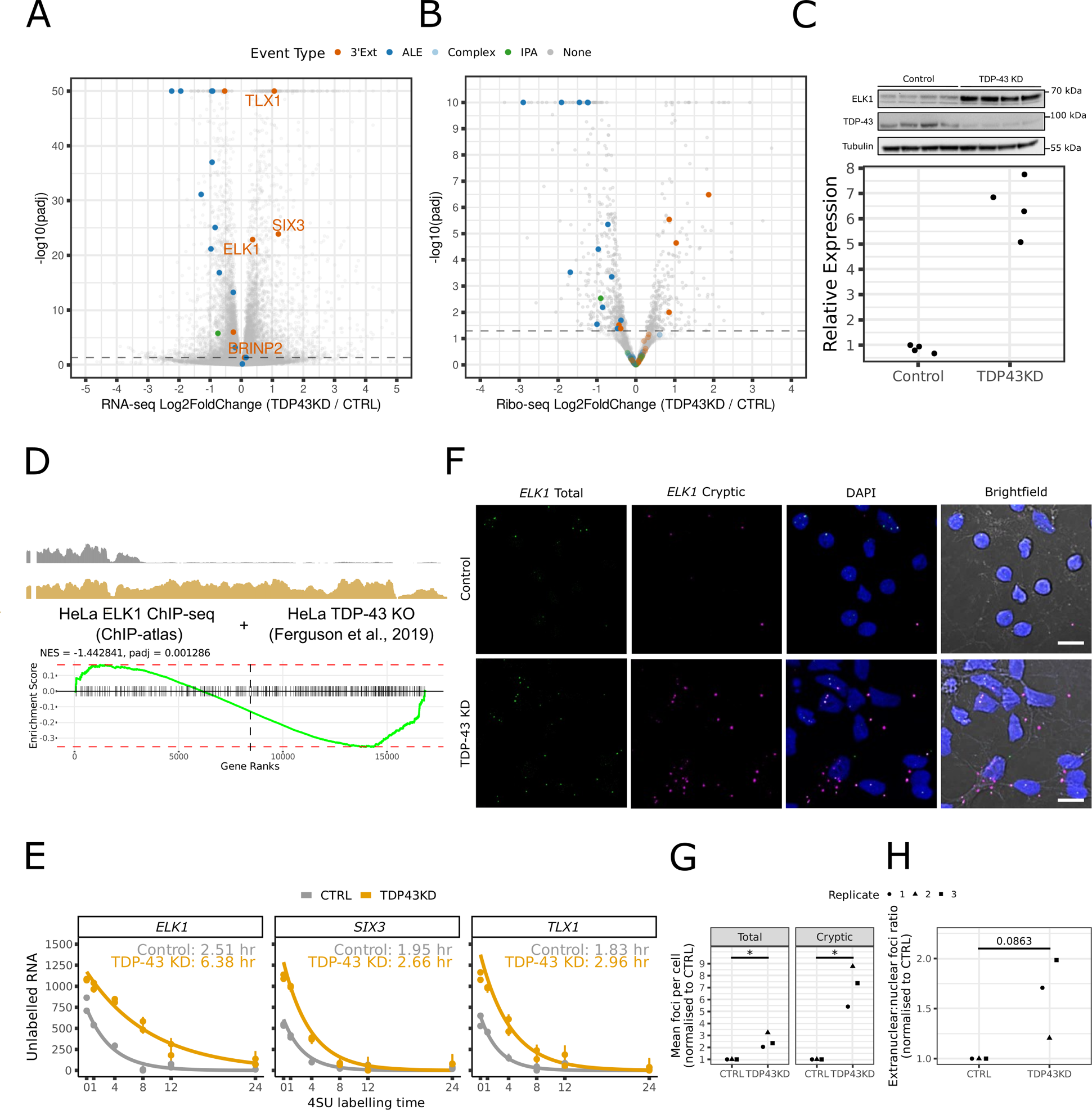
Cryptic 3’UTR extensions in transcription factor RNAs lead to increased RNA and protein levels by increased RNA stability and altered localisation. A) Volcano plot of differential expression analysis of RNA-seq data between TDP-43 knockdown (TDP43KD) and control (CTRL) i3Neurons. Cryptic 3’Ext genes with increased translation (Fig. 3B) are highlighted in orange. Genes with a −log_10_ transformed Benjamini-Hochberg adjusted p-value greater than 50 are collapsed to 50 for visualisation purposes. B) Volcano plot of differential expression analysis of Ribo-seq data between TDP-43 knockdown (TDP43KD) and control (CTRL) i3Neurons. Genes are highlighted if they contain a cryptic 3’Ext (orange), ALE (blue) or IPA (green) event. Genes with a −log_10_ transformed Benjamini-Hochberg adjusted p-value are collapsed to 10 for visualisation purposes. C) Western blot analysis of ELK1 protein levels in Halo-TDP-43 i3Neurons. (Top) Western blot showing increased ELK1 protein expression upon TDP-43 KD in Halo-TDP-43 i3Neurons. (Bottom) Quantification of ELK1 band intensities normalised to Tubulin in control (CTRL) and TDP-43 knockdown (TDP43KD) Halo-TDP-43 i3Neurons. D) Analysis of ELK1 transcription factor activity. (Top) Coverage trace in control (black) and TDP-43 knockout (gold) samples for the *ELK1* cryptic 3’Ext in HeLa cells^39^. (Bottom) Enrichment plot for ChIP-seq defined ELK1 target genes in TDP-43 knockout HeLa cells. The green line corresponds to GSEA’s running sum statistic, and red horizontal dashed lines mark the maximal enrichment score among upregulated and downregulated genes. The vertical dashed line demarcates the rank between upregulated and downregulated genes in the evaluated gene set. Black vertical bars correspond to locations of ELK1 target genes in the ranked gene set. The black text reports the normalised enrichment score (‘NES’, normalised to the mean enrichment score of random samples of the same size as the gene set) and Benjamini-Hochberg adjusted p-value (‘padj’) E) Decay curve for RNA produced before 4SU labelling in i3Neuron SLAM-seq data for control (grey) and knockdown (orange) samples. Solid curves indicate the fitted estimate of the level of old RNA for each condition. Individual samples are shown as points with the upper and lower 95% credible interval shown as error bars. GrandR-estimated half-lives for control (grey) and knockdown (orange) samples are reported in the inset text for each gene. F) Representative images for FISH probes targeting the annotated (‘*ELK1* Total’, green) 3’UTR and cryptic 3’UTR specific (*‘ELK1* Cryptic’, magenta) sequences of *ELK1* in control (top row) and TDP-43 knockdown (bottom row) i3Neurons. The white lines in the ‘Brightfield’ column represent scale bars (10 µm). G) Quantification of total FISH probe signal for the probes targeting the annotated 3’UTR (‘Total’) and cryptic 3’UTR specific (‘Cryptic’) sequences of *ELK1*. Different shapes represent independent replicates. Mean foci counts per cell (n=10 images) are normalised with respect to the control sample from the same replicate (Methods). A single asterisk (*) represents a Benjamini-Hochberg adjusted p-value < 0.05 from a one-sample t-test on log-transformed ratios (Methods). H) Subcellular quantification of FISH signal for probes targeting the annotated 3’UTR region (‘Total *ELK1*’) of *ELK1*. Different shapes represent independent replicates. The mean ratio of extranuclear:nuclei foci counts (n=10 images) is normalised with respect to the control sample from the same experimental replicate (Methods). The numeric label represents the Benjamini-Hochberg adjusted p-value from a one-sample t-test on log-transformed ratios (Methods).

Interestingly, the three 3’Ext-containing genes that were most upregulated at both RNA and translation levels (**Fig. 3A,3B**) encode for three transcription factors (TFs): *ELK1*, *SIX3,* and *TLX1*. The regulation of these 3’Ext events is reproducible across *in vitro* datasets (**Supplementary Fig. 1**). As *ELK1* increase has previously been associated with neuronal toxicity^33–35^ and its levels are consistently higher in mature neurons, compared to *SIX3* and *TLX1,* which are associated with neuronal development^36,37^, we decided to focus our investigations on *ELK1*. We tested whether the increase in Ribo-seq also corresponded to an upregulation of steady-state protein, and western blots confirmed a significant increase in ELK1 protein expression upon TDP-43 knockdown in i3Neurons (**Fig. 3C**).

We next asked whether the activity of ELK1, which functions as a TF in the ternary complex factor (TCF) family^38^, could be altered in the context of TDP-43 loss. We assessed whether ELK1 target genes defined by ChIP-seq in HeLa cells were also affected in TDP-43 knockout HeLa cells^39^, where the cryptic 3’Ext is robustly upregulated (**Fig. 3D**). Using GSEA, we observed a significant change in ELK1 target gene expression upon TDP-43 knockout (**Fig. 3D**). This suggests that cryptic 3’Exts can lead to change in function in the context of TDP-43 loss.

### TFs with cryptic 3’UTR Extensions have increased RNA stability and cytoplasmic RNA localisation

We investigated the mechanisms by which cryptic 3’UTRs could mediate increased translation levels of *ELK1*, *SIX3*, and *TLX1*. We revisited differential splicing analysis of i3Neuron RNA-seq datasets^10,13^ and confirmed that cryptic 3’Exts are the only differential RNA processing events occurring in these 3 TF RNAs upon TDP-43 depletion.

As alternative 3’UTRs have been linked to differences in RNA stability^40^, we reasoned that increased RNA stability could account for changes in overall RNA abundance and translation levels. To investigate changes in RNA stability in i3Neurons with TDP-43 depletion, we performed SLAM-seq^41^, which allows the detection of newly synthesised RNAs through incorporation of a uridine analogue (4sU). Different lengths of 4sU treatment allow to estimate gene-level RNA half lives. We observed increased half lives in cryptic 3’Ext containing genes *ELK1*, *TLX1,* and *SIX3* (**Fig. 3E**). This suggests that increased RNA abundance and translation of cryptic 3’Ext genes are mediated by increased RNA stability.

Given that translation depends on extra-nuclear localisation of mRNAs, we tested whether altered subcellular localisation of transcripts could be induced by 3’Ext and also contribute to the increased translation levels^42–45^. Focussing on the *ELK1* cryptic 3’Ext, we designed probes to recognise the common proximal sequence and the distal sequence specific to the 3’Ext, and performed fluorescent in-situ hybridisation (FISH) in i3Neurons (**Fig. 3F**). Consistent with RNA-sequencing, upon TDP-43 knockdown we observed a significant increase in total foci for both the total and cryptic-specific probes, with a more pronounced increase for the cryptic-specific probe (**Fig. 3G**). While there was a trend for elevated extranuclear localisation of *ELK1* (**Fig. 3H**), this increase did not reach significance. These findings suggest that increased RNA stability is likely the main driver of increased *ELK1* RNA abundance and translation.

## Discussion

Defining TDP-43 RNA targets is critical to understanding the molecular consequences of nuclear TDP-43 depletion. To date, efforts have mainly focussed on the consequences of altered splicing and have successfully identified key targets that are being pursued as therapeutic targets and potential biomarkers for TDP-43 pathology^10,11,14,46,47^. Although TDP-43 is involved in multiple aspects of RNA processing, including polyadenylation^4–6^, this has been largely understudied due to the lack of effective tools to address these questions. Here, we develop a pipeline to detect and quantify novel APA events from total RNA-seq and apply it to a wide range of neuronal TDP-43 loss of function datasets to define cryptic APAs, a novel category of cryptic RNA processing events of potential relevance to ALS/FTD. iCLIP and TDP-43 binding motif analyses support a direct regulation of these events by TDP-43, in which TDP-43 loss can both weaken conventional polyA sites and derepress cryptic APA. Similarly to splicing, where TDP-43 can both repress or enhance exon inclusion, TDP-43 can therefore have a dual action on transcript termination. Importantly for disease relevance, and similarly to cryptic splicing, numerous cryptic APA events can be detected in post-mortem tissue and are specifically expressed upon TDP-43 pathology.

When we then moved to investigate the impact of cryptic APAs on RNA levels and translation, and found that IPAs and ALEs had either no impact or induced a reduction of transcript levels in RNA-seq and Ribo-seq analyses - in line with previous observations on known cryptic ALEs as *STMN2*^46,47^. Recent work has demonstrated that cryptic exon-containing transcripts can be translated and produce cryptic peptides that could serve as biomarkers of TDP-43 pathology^13,14^. As cryptic ALE and IPA events are mostly predicted to be insensitive to nonsense mediated decay, they are likely to give rise to cryptic peptides; e.g. cryptic ALE *RSF1* encodes a cryptic peptide that is detected in CSF of ALS patients^13^. Previous work has identified cryptic ALEs as their novel splice junction can be detected by numerous splice-detection packages^13,27,48^. Conversely, IPAs have been harder to identify and further work should consider whether these cryptic IPA events can be detected in patient brains and biofluids as an indirect measure of TDP-43 pathology.

Surprisingly 3’Ext events in the three transcription factors ELK1, SIX3, and TLX1 were associated with transcript upregulation, increased translation and protein levels. We found this to be associated with an increase in RNA stability leading to an increase in cytoplasmic localisation. Thus, in contrast to the conventional model of TDP-43 regulated cryptic splicing leading to reduced protein levels or to altered proteins containing cryptic peptides, cryptic 3’Ext can be associated with increased protein levels, outlining a novel consequence of TDP-43 cryptic RNA processing.

*ELK1*, *SIX3,* and *TLX1* 3’Ext are reliably induced upon TDP-43 depletion across our *in vitro* datasets, suggesting they are not cell-type specific, sensitive TDP-43 targets. These three TFs have been studied in the neuronal context, although *SIX3* and *TLX1* are primarily expressed in the developmental stage^36,37^. Our work therefore focused on *ELK1* and we were able to utilise HeLa cell data in order to show that TDP-43 loss can induce changes in *ELK1* target genes. *ELK1* promotes axonal outgrowth^49^ and is increased in Huntington’s disease models where it can have a neuroprotective role^50^. *ELK1* overexpression has also been linked with neurotoxicity through interaction with components of the mitochondrial permeability-transition pore complex^34^, and dendrite-specific overexpression of *ELK1* mRNA induced cell death in a transcription- and translation-dependent manner^33^, supporting a potential contribution of this cryptic APA to pathogenesis. Further work is needed to investigate the functional relevance of increased *ELK1*, *SIX3*, and *TLX1* expression in models of TDP-43 proteinopathy.

We focussed on identifying cryptic APA events, as their extreme expression changes upon TDP-43 loss renders them favourable therapeutic and biomarker targets. Accompanying manuscripts by Arnold et al. and Zeng et al. investigate APA dysregulation more generally upon TDP-43 loss and show it is widespread, in accordance with our findings in **Fig. 1B**, can occur in ALS-FTD related genes (Zeng et al., co-submitted) and can lead to change in function (Arnold et al., co-submitted), underscoring the potential relevance of APA in disease pathogenesis. We note that several targets (e.g. *CNPY3, ELK1, ARHGAP32)* are commonly identified across the studies despite diverging methodological approaches, underlying the consistency of our observations. Importantly, similarly to our findings for *ELK1*, *SIX3*, and *TLX1*, both Zeng et al. and Arnold et al. manuscripts also find that APAs can lead to upregulation of normal protein levels, consolidating this a general consequence of TDP-43 loss. Our studies collectively demonstrate that dysregulated APA is a general consequence of nuclear TDP-43 loss in ALS-FTD.

In summary, we provide a compendium of cryptic APA events determined by TDP-43 loss as a resource for studying RNA dysregulation and identifying novel biomarkers in ALS. Our work also shows that cryptic RNA processing can lead to an increase in protein expression and function, expanding the molecular consequences of TDP-43 loss and pathology, with implications for disease pathogenesis and therapeutic target identification.

## Supporting information

NYGC ALS Consortium Author List

## Methods

### CRISPRi knockdown in human iPS cells, and differentiation and culture of i3Neurons

CRISPRi knockdown experiments were performed in the WTC11 iPSC line harbouring stable TO-NGN2 and dCas9-BFP-KRAB cassettes at safe harbour loci^51^. CRISPRi knockdown of TDP-43 in iPSCs was achieved using sgRNA targeting the transcription start site of TARDBP (or non-targeting control sgRNA)^10^, delivered by lentiviral transduction. sgRNA sequences were as follows: non-targeting control GTCCACCCTTATCTAGGCTA; and TARDBP GGGAAGTCAGCCGTGAGACC. iPS cells were differentiated into cortical-like i3Neurons as described previously^10,52^ and fixed 9 days after re-plating for RNA-fluorescence in-situ hybridisation (FISH). For RNA-seq experiments (‘Humphrey i3 cortical’), i3Neurons were induced as previously described^52^ with the addition of SMAD and WNT inhibitors^53^ (SB431542 10 µM; LDN-193189 100 nM; XAV939 2 µM all from Cambridge Biosciences). After induction, cells were cultured in BrainPhys Media (StemCell Technologies) with 20 ng/ml BDNF (PeproTech), 20 ng/ml GDNF (PeproTech), 1x N2 supplement (Thermo Fisher), 1 x B27 supplement (Thermo Fisher), 200 nM Ascorbic Acid (Sigma), 1 mM dibutyryl cyclic-AMP (Sigma) and 1 µg/ml Laminin (Thermo Fisher) as previously described^54^ and harvested 30 days after differentiation.

An iPSC line with a N-terminal HaloTag on both endogenous copies of TDP-43 (Halo-TDP-43 i3Neurons) was generated by CRISPR/Cas12 gene editing. The parental cell line used was the WTC11 cell line with integrated dCas9-Krab and NGN2 cassettes as mentioned previously^51^. The HDR template used was addgene plasmid 178131. Editing was done with Cas12 crRNA (Integrated DNA Technologies) with GGAAAAGTAAAAGATGTCTGAAT as the targeting sequence. Recombinant Cas12 (Cpf1 ultra, Integrated DNA Technologies) was electroporated with HDR template and Cas12 crRNA using the P3 Primary Cell 4-D Nucleofector kit (V4XP-3024 Amaxa). iPSCs were then single cell plated and positive colonies were selected with HaloTag TMR dye (Promega) and verified by PCR of genomic DNA.

For PROTAC mediated knockdown of Halo-TDP-43, i3Neurons were treated with HaloPROTAC-E^55^ (30 nM) on DIV14 and harvested on DIV28.

Strand-specific, poly(A) enriched sequencing libraries for the ‘Humphrey i3 cortical’ dataset were prepared using the Kapa mRNA Hyper Prep kit. 100ng total RNA was used as input material for poly(A)+ mRNA capture. Fragmentation was performed for 6 minutes at 85°C to obtain a target fragment size of 300-400bp, and 13 cycles of PCR amplification were performed. The resulting libraries were sequenced 2×150bp on the Illumina NextSeq2000 machine. Samples were processed as previously described^56^ using the RAPiD-nf nextflow pipeline. Briefly, adapters were trimmed from raw reads using Trimmomatic^57^ v0.36, and reads were aligned to the GRCh38 genome build using gene models from GENCODE v30^58^ with STAR^59^ v2.7.2a. The RAPiD-nf pipeline is available at https://github.com/CommonMindConsortium/RAPiD-nf/.

### Fluorescent in-situ hybridisation (FISH)

Cortical-like i3Neurons were cultured on 13-mm glass coverslips and fixed in 4% PFA/sucrose on day 9. RNA-FISH was performed using the QuantiGene ViewRNA ISH Cell Assay kit (Invitrogen, QVC0001), according to manufacturer’s instructions. Protease was used at 1:1000 dilution. Two probe sets were used to detect the canonical *ELK1* transcript (TYPE 4 probe, 488-nm) or specifically the distal 3’UTR cryptic extension (TYPE 1 probe, 550-nm). Confocal images were acquired with a LSM980 laser-scanning confocal microscope with Airyscan 2 (Zeiss), using a 40X oil immersion objective.

For each biological replicate, ten images were acquired for the control and TDP-43 knockdown conditions. For each image, foci for both probes were counted within the 106.07 microns by 106.07 microns field of view on Fiji/ImageJ using the maximum intensity Z-projection function to flatten the 2-µm-thick Z-stack. The Find Maxima function using the same prominence setting between conditions was performed to quantify total numbers of RNA foci. To separately count nuclear and cytoplasmic foci, the Cell Counter plugin was used. For each probe and field of view, the total number of foci was divided by the number of DAPI-stained nuclei to give the average number of foci/cell. To calculate the nuclear:extra-nuclear ratio for the ‘Total *ELK1*’ probe, the number of nuclear foci was divided by the number of extra-nuclear foci in each field of view. For each probe and condition, the mean number of foci/cell and nuclear:extra-nuclear ratio was calculated from the ten images and normalised, for each biological replicate, to the respective control condition. Statistical significance was evaluated using a one sample t-test with a logarithmic transformation and the Benjamini and Hochberg false discovery rate procedure, testing the null hypothesis that mean = log(1).

### Western blots

Halo-TDP-43 i3Neurons were homogenised in lysis buffer (25 mM Tris-HCl, 150 mM NaCl, 1% NP-40, 1% Glycerol, 2 mM EDTA, 0,1% SDS, protease inhibitor (cOmplete^TM^ EDTA-free protease inhibitor cocktail, Roche), phosphatase inhibitor (PhoSTOP^TM^, Roche)). Samples were loaded on a NuPAGE 4-12% Bis-Tris protein gel (Invitrogen), which was run in NuPAGE MOPS buffer. Proteins were transferred onto PVDF blotting membrane (Amersham), through wet transfer for 1h 30 min at 200 mA in transfer buffer (25 mM Tris, 192 mM glycine, 20% methanol). The membrane was blocked in 5% milk in TBST (20 mM Tris, 150 mM NaCl, 0.1% Tween-20) and incubated overnight with primary antibodies diluted in 5% milk in TBST (anti-ELK1 (Abcam ab32106) 1:500, anti-TDP-43 (Abcam, ab104223) 1:2000, anti-tubulin (Sigma-Aldrich, MAB1637) 1:5000). After 1h-incubation with HRP-conjugated secondary antibodies diluted in 5% milk in TBST (anti-mouse HRP (BioRad, 1706516) 1:10000, anti-rabbit HRP (BioRad 1706515) 1:10000), the membrane was developed using Immobilon Classico HRP substrate (Sigma) and the Bio-Rad ChemiDoc system.

### SH-SY5Y & SK-N-BE(2) TDP-43 KD and sequencing

SH-SY5Y and SK-N-BE(2) cells were transduced with a SmartVector lentivirus (V3IHSHEG_6494503) containing a doxycycline-inducible shRNA cassette for TDP-43. Transduced cells were selected with puromycin (1 μg/ml) for one week, before being plated as single cells and expanded to obtain a clonal population. Cells were grown in DMEM/F12 + Glutamax (Thermo) supplemented with 10% FBS (Thermo) and 1% PenStrep (Thermo).

For induction of shRNA against TDP-43 cells were treated with the following amounts of doxycycline hyclate (Sigma), and collected after 10 days:

- For experiments in SH-SY5Y cells (curves), 75 ng/ml.
- For experiments in SH-SY5Y cells (CHX), 25 ng/ml.
- For experiments in SK-N-BE(2) cells, 1000 ng/ml.

RNA was extracted from SH-SY5Y and SK-N-BE(2) cells using the RNeasy mini kit (Qiagen) following the manufacturer’s protocol including the on-column DNA digestion step. RNA concentrations were measured by Nanodrop and 1,000 ng of RNA was used for reverse transcription. Samples undergoing RNA sequencing were furthermore assessed for RNA quality on a TapeStation 4200 (Agilent), resulting in RNA integrity number (RIN) above 9.4 for all samples.

Sequencing libraries were prepared with polyA enrichment using a TruSeq Stranded mRNA Prep Kit (Illumina) and sequenced on an Illumina HiSeq 2500 or NovaSeq 6000 machine at UCL Genomics with the following specifics:

- SH-SY5Y cells: 2×100 bp, depth > 40M/sample
- SK-N-BE(2) cells: 2×150 bp, depth > 40M/sample

### RNA-seq data processing

The ‘Brown’ SH-SY-5Y, SK-N-BE(2) and i3Neuron datasets were processed as previously described^10^. Unless otherwise stated, all short-read RNA-seq datasets were processed using the following pipeline. Raw reads in FASTQ format were quality trimmed for a minimum Phred score of 10 and otherwise default parameters using fastp^60^ (v0.20.1). Quality trimmed reads were aligned to the GRCh38 genome build using gene models from GENCODE v40^58^ with STAR^59^ (v2.7.8a). Quality trimmed reads are used as input for any tools that require FASTQ files as input (e.g. PAPA, Salmon). Our alignment pipeline is implemented in Snakemake^61^ and is available at https://github.com/frattalab/rna_seq_snakemake.

### SLAM-seq

SLAM-sequencing was performed on cortical-like i3Neurons following protocols adapted from Herzog et al.^41^. Samples were treated with 100 µM 4sU on Day 7 for 0, 1, 4, 8, 12, 24 hours before immediate wash with phosphate buffered saline (PBS). Each time point had 2 replicates for both control and TDP-43 knockdown excluding 4 hours where one of the control replicates did not pass RNA quality controls and so was not submitted for sequencing.

RNA was extracted using the Qiagen RNA isolation and purification kit. RNA concentration was estimated using a Nanodrop Microvolume Spectrophotometer (Thermofisher). After ensuring an adequate amount of RNA in each sample, iodoacetamide (IAA) treatment was applied to each, facilitating the thiol modification of incorporated 4sU.

Sequencing libraries were prepared with Kapa RiboErase RNA Hyper kit and sequenced (2 × 250 bp) on an Illumina NovaSeq SP. Using the ‘rna_seq_snakemake’ alignment pipeline (https://github.com/frattalab/rna_seq_snakemake), raw FASTQ files were quality trimmed using fastp^60^ with the parameter “qualified_quality_phred: 10”, and aligned without soft-clipping to the GRCh38 genome build using STAR^59^ (v2.7.0f) with gene models from GENCODE v34^58^. GRAND-SLAM (v2.0.7b) was run on the aligned data using gene models from GENCODE v34^58^ using the “-trim5p 10 -trim3p 10” parameter to ignore mismatches at the ends of reads. The output files containing the estimated new-to-total RNA ratios (NTR) of each gene were used to estimate the half-life of each gene using the recommended workflow in grandR^62^.

### PAPA - pipeline to detect cryptic last exons

Whilst there are many tools for de-novo detection of alternative polyadenylation events within 3’UTRs events from RNA-seq data, all of them suffer from poor performance with respect to matched 3’end sequencing approaches^63,64^. Few tools have focused on the detection of gene-body internal poly(A) sites and definition of the complete last exon structure, which is essential if one aims to predict putative encoded peptides. Aptardi is a deep-learning based approach to refine predicted 3’ends of reference or assembled transcriptomes^65^, but a benchmarking study found the compute times and resources to be unviable for performance evaluation^64^. TECTool trains a machine-learning model on annotated last exons using transcriptomic features to classify novel intronic last exons defined upstream of polyA sites from the PolyASite atlas^66^. However, as of v0.4 it can only define ALEs and only supports single-end RNA-seq data, which would substantially impact sensitivity to detect events.

General purpose bulk RNA-seq transcript assemblers such as StringTie^19^ could be used to identify all alternative novel last exon structures, but they show poor performance in defining precise terminal exon boundaries^66^. Previous benchmarking attempts have suggested that transcript assemblers exhibit poor performance at defining full transcript structures, but much better performance at defining individual exons^67^. This suggests that a scalable solution could be to refine the 3’end predictions of exons predicted by transcript assemblers.

Broadly inspired by a previous workflow combining matched short read and 3’ enriched sequencing^18^, our approach is to extract last exons from StringTie assembled transcripts and filter based on proximity to 3’end-seq derived PolyA site annotations and/or presence of polyA signal sequences in the terminal predicted region. PolyA signal hexamer presence previously emerged as one of the most important features in discriminating intergenic expressed regions as 3’UTRs from other transcriptomic regions^68^.

### Pipeline setup

Transcript assemblies for individual samples are generated using StringTie v 2.1.7 (default settings) in annotation-guided mode. Individual sample assemblies are grouped according to their experimental condition and merged into a redundant assembly using Gffcompare^69^ v0.11.2. Condition-wise mean TPMs are calculated for each transcript, assigning a TPM of 0 if a transcript was not assembled in a particular sample. Transcripts are subsequently filtered for a minimum mean expression > 1 TPM, on the basis of a previous study which proposed such a filter to improve global accuracy of assembled transcripts with respect to matched long-read sequencing^70^.

Last exons are extracted from expression-filtered transcripts in a sample-wise manner using a custom script. Novel last exons are extracted using the following criteria:

- All predicted 3’ends must not overlap with any annotated exon
- The last intron of ALEs must be contained within an annotated intron and its 5’ss must exactly match an annotated 5’ss
- The last intron of putative events must overlap an annotated exon, with the predicted 5’end of the last exon exactly matching the 5’end of the overlapping annotated internal exon. If the putative last exon overlaps an annotated first exon, the matching of 5’ends is permitted a 100nt ‘slack’ to permit last exon capture despite imprecision of the predicted transcript 5’end, which is of secondary importance for our purposes and is a known limitation of transcript assembly tools with short-read RNA-sequencing
- Last exons overlapping annotated last exons must extend the most distal annotated last exon at a locus and exactly match at the 5’end of the annotated exon
- All ‘extension’ events must extend a known exon by a user-specified minimum distance (default 100nt)

Putative novel last exons are subsequently merged condition-wise into single GTF using a custom script. Filtering and refining of putative last exons for 3’end precision is subsequently performed condition-wise, with the aim of selecting a single representative last exon prediction for a given condition. Firstly, the distance (in either direction) from the 3’end of last exons to the nearest locus reported in the PolyASite 2.0 database^20^ is calculated. Any distance below a user-specified distance (default 100nt) is considered a match and retained for downstream analysis. Considering that 3’seq protocols on which the PolyASite database is based can provide nucleotide resolution of poly(A) sites, the 3’ends of matching last exons are updated to the matching site reported in the PolyASite database.

Given that the PolyASite database (as of version 2.0) lacks datasets with TDP-43 depletion and has limited neuronal cell datasets, it is possible it has incomplete coverage of polyadenylation sites specific to TDP-43 depletion and/or neuronal cell contexts. PolyA sites are characterised by an enriched nucleotide distribution around cleavage sites. Most notably, polyA signal hexamers, of which 18 variants exist^21^, are enriched approximately 21 nt upstream of cleavage sites. To rescue 3’ends of last exons that may not be represented in the PolyASite database, the final 100nt of putative last exon sequences are extracted and last exons are retained if an exact match to any of the previously defined 18 polyA signal hexamers^21^ is found. If a locus contains multiple predicted last exons, the last exon with a polyA signal hexamer located closest to the expected 21nt upstream distance is selected for each experimental condition.

Next, a combined transcriptome reference of novel and annotated last exons is generated. All last exons passing either the PolyASite or motif filter are retained for downstream analysis. Last exons of all annotated transcripts are extracted using a custom script. Last exons of each gene are assigned a common ‘last exon isoform identifier’ based on any overlapping sequence. 3’UTR extensions are assigned a distinct identifier, and the annotated last exon(s) they extend are grouped into a single identifier. This has the effect of comparing the usage of 3’UTR extension to all other annotated last exons of the same gene. In order to prevent misattribution of reads to internal last exon isoforms, any regions overlapping annotated first or internal exons are removed (i.e. only ‘unique’ regions of last exons are retained). Any last exons with a 3’end overlapping annotated first/internal exons are also completely removed from downstream analysis to produce a final reference of last exon isoforms for quantification.

Transcript sequences are extracted using gffread^69^ version 0.12.1 and used to produce a decoy aware transcriptome index constructed using Salmon^22^ (version 1.5.2) with full genome sequence (Grch38 build) used as decoys^71^. Samples are subsequently quantified against the last exon reference using Salmon^22^ v1.5.2 with the ‘--gcBias’ & ‘--seqBias’ flags. Transcript per million (TPM) values of individual transcripts are summed according to their assigned last exon isoform ID, and estimated counts are generated using the ‘countsFromAbundance=lengthScaledTPM option’ in the tximport^72^ package (version 1.26.0). The counts matrix is optionally used as input to DEXSeq^23^ v1.44.0 to test for differential isoform usage between experimental conditions. A relative polyA site usage is further calculated for each gene by dividing the expression (in TPM units) of each last exon isoform by the sum of expression of all isoforms of the gene.

PAPA v0.2.0 is implemented as a Snakemake^61^ pipeline. All interval operations in Python are performed using the PyRanges^73^ package, and genomic sequence operations using a combination of pyfaidx^74^ and BioPython^75^. Package dependencies are managed using conda environments, including an ‘execution’ environment with a minimal Snakemake installation. The code is available at https://github.com/frattalab/PAPA.

### Identification of cryptic last exons with PAPA

To generate a common transcript reference against which to quantify last exons, we first ran PAPA in ‘identification’ mode to predict novel last exons across all stranded datasets in our compendium (All i3Neuron datasets, two of the SH-SY5Y datasets and one SK-N-BE(2) dataset). StringTie^19^ transcript assembly was performed in an ‘annotation-guided’ manner using a filtered GENCODE v40^58^ human transcriptome reference. The reference was filtered first for transcript models with a ‘transcript support level’ tag value of at least 3 that belong to protein-coding or lncRNA genes. Transcript models with the ‘mRNA_end_NF’ tag, denoting transcripts with unsupported 3’ends, are also removed as performed by LABRAT^76^.

Gene transfer format (GTF) files of predicted last exons across all datasets were subsequently combined into a single GTF of novel last exons using the custom script ‘combine_novel_last_exons.py’ script available in the PAPA repository. All datasets were then quantified and assessed for differential usage using a unified transcriptome reference of combined novel last exons and annotated last exons (from the same filtered GTF used for transcript assembly). All differential usage tests were performed using the standard DEXSeq workflow without additional covariates, with the exception of the Klim et al i3 motor neuron dataset^47^ where the date of differentiation was added as a covariate. Cryptic last exons were defined as isoforms with an Benjamini-Hochberg adjusted p-value of < 0.05 from DEXSeq, mean usage in control samples < 10 % and increase in mean usage in knockdown samples > 10 %.

Following manual inspection of cryptic events we observed frequent IPA calls resulting from regions with intron retention, where the predicted 3’end can be attributed to regions with marked well of reduced coverage (likely due to repetitive sequence) but similar levels of coverage either side of the well and at the intron-exon boundaries. We therefore manually curated cryptic IPA events to mitigate these artefacts. We do not anticipate intronic ALEs to be similarly affected because their 5’end is defined by a novel splice junction.

### TDP-43 iCLIP analysis

The SH-SY5Y TDP-43 iCLIP data was generated and processed as previously described^10^, and the raw data is available at E-MTAB-11243. iCLIP peaks from the two independent replicates were merged into non-redundant intervals for all subsequent analysis.

Cryptic events are defined as last exon isoforms passing cryptic thresholds in any *in vitro* dataset. The probability of detecting TDP-43 binding events via iCLIP is influenced by the abundance of target RNAs, but by pooling cryptic events across datasets we cannot control for the confounding influence of RNA expression between groups. We therefore defined background events as isoforms that were assessed for differential usage (see above criteria) in all SH-SY5Y datasets and had a padj > 0.05 across all datasets. Given that the cryptic group contains events identified across all datasets (with no guarantees of expression in SH-SY5Y datasets), we therefore penalise the detection of TDP-43 binding in the cryptic group and bias against observing enriched binding in this group.

To define representative intervals for 3’Ext events, the most distal annotated polyA site is selected to represent the proximal site, and background events represent loci with a predicted novel 3’UTR extension. For other event categories, background events include annotated and novel events. However, our approach to define a common last exon reference across datasets and defining last exon isoforms (see above) can result in non-redundant intervals being predicted for the same last exon isoform. As such, we implemented a collapsing strategy to define a single representative interval for each event. First, overlapping novel predictions are filtered for those that match a site from the PolyASite atlas. If distinct reference sites are reported for the same isoform, the site that is predicted in the most independent datasets is selected as representative. If distinct sites are detected in the same number of independent datasets, the more proximal site is arbitrarily selected as representative. Finally, because the standard PolyASite atlas cluster intervals were used for matching, distinct 3’end predictions can overlap with cluster intervals. In these cases the site that is closest to the PolyASite representative coordinate is selected. If these distinct cluster-overlapping sites are equidistant from the representative coordinate, the most distal coordinate is arbitrarily selected. If no isoforms match an atlas site (i.e. contains a polyA signal sequence), we first attempt to select a representative site whose putative polyA signal motif minimises the deviance from the characteristic position 21nt upstream of the polyA site. If multiple motifs are equidistant from the expected position, the most proximal site is arbitrarily selected as representative. For ALE and IPA events, we noted that collapsing at the 3’end still resulted in distinct intervals for each last exon. 9 background IPA events still had distinct 3’end coordinates following the filtering criteria above, and the interval with the most distal 3’end was arbitrarily selected as the representative interval. 4 background ALE events had distinct 5’end coordinates following the above filtering, and the most proximal 5’end (i.e. shortest exon) is arbitrarily selected as the representative interval. Given the very small proportion of events affected by these arbitrary criteria, we do not anticipate they will substantially affect the analysis.

To construct metaprofiles of TDP-43 binding, Single nucleotide regions representing the genomic landmarks were extended by 500nt in both directions and per-position coverage by iCLIP peaks was then calculated, assigning a value of 1 when a given position overlaps an iCLIP peak. All interval operations were performed using bedtools^77^ version 2.31.0. The mean (equivalent to the fraction of events that have an overlapping iCLIP peak) and standard error of coverage is then calculated for each position relative to the landmark. Confidence intervals around the mean coverage in the maps correspond to +/− 1 standard error. Both mean coverage and confidence intervals are visualised following LOESS smoothing with the ‘span’ parameter set to 0.1.

### De-novo motif enrichment analysis

To perform de-novo motif enrichment, we adapted PEKA^24^, which identifies kmers with positional enrichment at iCLIP peaks relative to background cross-link sites whilst normalising to the general occurrence in the surrounding genomic context. Therefore, we can substitute iCLIP peaks and global cross-link sites for cryptic and background landmarks respectively to identify positionally enriched kmers with respect to cryptic landmarks. For all comparisons, we ran PEKA to search for enriched 6-mers in the proximal window of interest set to 250nt (the broad window in which iCLIP peaks were observed) and the distal window set to 500nt (to maintain consistency with the overall search space for iCLIP peaks). The ‘percentile’ flag was set to 0 to switch off thresholding of background regions based on read count, and the ‘relpos’ flag to 0 to consider all positions in the proximal window when calculating the enrichment score.

Preferred TDP-43 binding 6-mers were extracted from Halleger et al.^6^. Briefly, The 6-mers were defined using PEKA as the as the top 20 most enriched kmers around intronic iCLIP crosslinks across all WT, A326P, G294A, G335A, M337P, Q331K and a 316del346 GFP-TDP-43 in HEK293 cells. The 20 were subsequently separated into the following three groups based on a gradient of enrichment in WT and G335A TDP-43 with respect to A326 and 316del346 variants and their consensus sequence:

- YG-containing [UG]n 6-mers - UGUGUG, GUGUGU, UGUGCG, UGCGUG, CGUGUG, GUGUGC
- YA-containing [UG]n 6-mers - AUGUGU, GUAUGU, GUGUAU, UGUGUA, UGUAUG, UGCAUG
- AA-containing [UG]n 6-mers - GUGUGA, AAUGAA, GAAUGA, UGAAUG, AUGAAU, GUGAAU, GAAUGU, UUGAAU

Where ‘Y’ corresponds to a pyrimidine nucleotide. To assess their over-representation among enriched 6-mers relative to cryptic landmarks, we performed a one-sided gene-set enrichment analysis (GSEA) using fgsea^32^ version 1.24.0 with default settings for each cryptic landmark. The three 6-mer groups and the union of all three groups were provided as input pathways, and kmers were ranked by their PEKA score. After independent runs for each landmark, Benjamini-Hochberg adjusted p-values were calculated with respect to the all tested landmarks and 6-mer sets and used to evaluate statistical significance.

To generate maps of coverage of specific kmers, we used cv_coverage^78^ v1.1.0 (https://github.com/ulelab/cv_coverage) to scan for occurrences of the YG-containing [UG]n 6-mers in a 500nt window around cryptic and background landmarks, disabling weighting the occurrence by cDNA count. For coverage plots, the percentage occurrences of each 6-mer were summed separately for the cryptic and background regions. The percentage occurrences were converted to mean coverages and visualised as described for iCLIP maps.

The adapted PEKA code is available at the ‘output_mods’ branch of the following forked copy of the PEKA repository https://github.com/SamBryce-Smith/peka. A snakemake pipeline to run PEKA and cv_coverage is available in the ‘motifs/peka_snakemake’ directory of the ‘tdp43-apa’ repository.

### Post-mortem RNA-seq analysis

#### FACS-seq data processing

Sequenced reads from FACS-sorted frontal cortex neuronal nuclei^26^ and were processed as described in Brown et al^10^. The data are available on the Gene Expression Omnibus at GSE126543.

### Quantification of cryptic last exons in post-mortem FACS-seq data

Nuclear RNA-seq libraries contain both nascent and processed RNA. We therefore constructed decoy transcript models that include intron retention at the ALE and IPA loci to limit the confounding effect of nascent RNAs on transcript quantification^22^. Cryptic last exons are first classified as ALE, IPA or 3’Exts using the same criteria as PAPA, and decoy transcript models are subsequently generated separately for each event type.

For IPA events, the unique cryptic last exon sequence is extended to incorporate the annotated internal exon (up to its 5’ boundary). Then, a ‘spliced’ decoy transcript is generated that traverses the annotated internal exon to the downstream exon for all annotated transcripts, and an ‘intron retention’ decoy transcript is generated that contains the same pairs of internal exons and the intervening intron. For ALEs, a ‘retained intron’ decoy transcript is generated that consumes the complete intronic sequence in which the last exon is contained. No decoy transcript models are generated for 3’Ext events. Decoy transcript identifiers are appended with suffixes to differentiate from cryptic ALEs and last exon identifiers are generated (excluding decoys) with respect to novel and annotated last exons as in PAPA to allow calculation of % PAU usage. The script used to generate the decoy-augmented last exon reference is available at ‘add_decoys_to_gtf.py’ in the ‘tdp43-apa’ GitHub repository.

The decoy-augmented reference quantified with Salmon v1.8.0^22^ using the ‘salmon’ sub-pipeline available at https://github.com/frattalab/rna_seq_single_steps. As with PAPA, samples are quantified against a decoy-aware transcriptome index with full genome sequence (GRCh38 build) used as decoys^71^ and the ‘--gcBias’ and ‘--seqBias’ flags enabled.

Calculation of % polyA usage is performed using a copy of the ‘tx_to_polyA_quant.R’ script from the PAPA repository. Sample-wise differences in % polyadenylation site usage is calculated by subtracting the usage in the TDP-43 positive population from the TDP-43 negative population, such that a positive difference indicates the cryptic APA has a higher relative expression in the population with TDP-43 depletion.

### New York Genome Centre (NYGC) RNA-seq data

The sequencing libraries were generated^27,79^ and processed^13^ as previously described. Samples were classified into disease subtypes as previously described^13^. Briefly, FTD subtypes were classified by pathology according to the presence of TDP-43 inclusions (FTLD-TDP), FUS or Tau aggregates. ALS patients were sub-categorised based on presence (ALS-non-TDP) or absence (ALS-TDP) of reported SOD1 or FUS mutations. The following samples were considered as regions where TDP-43 pathology (and specific cryptic junction expression) is expected; motor (ALS-TDP), frontal and temporal cortex samples (FTLD-TDP, ALS-TDP), cervical, lumbar and thoracic spinal cord samples (ALS-TDP).

We opted to quantify ALE events using junction reads, which provide direct quantification of the occurrence of a splicing event. As of version 0.2 PAPA does not directly report splice junctions associated with ALE events. However, as the filtering criteria applied by PAPA requires putative ALE events to have a terminal splice junction with a direct match to an annotated 5’ss, it is possible to infer splice junctions from reference annotation using just the reported last exon coordinates. For ALEs fully contained within annotated introns, the splice junction is defined from the intron start to the start of the ALE. If last exons are distal to the annotated gene, then the closest upstream annotated intron is found. The splice junction is subsequently defined as the region from intron start to the start of the ALE. Finally, for annotated ALEs all annotated introns that terminate at the ALE are reported as splice junctions for the event. The above steps are implemented in a custom script ‘last_exons_to_sj.py’ available at the ‘tdp43-apa’ GitHub repository.

Splice junctions for cryptic ALEs and cryptic splice junctions identified in cortical-like i3Neurons^13^ were quantified across the NYGC RNA-seq cohort by extracting counts for provided junctions from the ‘.SJ.out.tab’ files produced by STAR^59^. The code is implemented in the ‘bedops_parse_star_junctions’ v0.1.0 Snakemake pipeline and is available at https://github.com/SamBryce-Smith/bedops_parse_star_junctions.

We defined detection criteria to prioritise cryptic splice junctions that are specifically in tissue types and samples with expected TDP-43 pathology. Junctions are considered expressed if at least two spliced reads are detected in a sample. Junctions are considered selectively expressed if expressed in at most 0.5 % of all samples where TDP-43 pathology is not expected and at least 1 % of samples where TDP-43 pathology is expected. We note that such criteria will exclude events with enriched expression in tissues with expected TDP-43 proteinopathy, but that have basal expression in unknown cell types not represented in our *in vitro* compendium. Such events may still have relevance in mechanisms of disease in specific cell types, but are less suitable for discriminating samples with TDP-43 proteinopathy.

### Ribo-seq analysis

i3Neuron Ribo-seq data was generated and processed as previously described^13^. Uniquely mapped reads were assigned to genes based on the union of annotated ‘CDS’ entries in the Gencode v34 standard annotation released using featureCounts^80^ version 2.0.1. Differential expression between TDP-43 knockdown and control was performed using DESeq2^81^ v1.38.3, and differentially translated genes were defined based on a Benjamini-Hochberg adjusted p-value threshold of 0.05. Any last exon passing our cryptic criteria in at least one of the i3 Neuron datasets (Brown i3 cortical, Seddighi i3 cortical, Humphrey i3 cortical) was considered for intersection with differentially translated genes.

Gene-set enrichment analysis was performed using fgsea^32^ version 1.24.0 with default settings. Cryptic 3’Ext, IPA & ALE containing genes were provided as input pathways, and moderated fold changes calculated with ‘lfcShrink’ function from DESeq2 package using the default apeglm^82^ method as the shrinkage estimator to rank genes. A threshold of 0.05 Benjamini-Hochberg adjusted p-value was used to determine statistical significance.

Read counting was performed using the ‘feature_counts’ sub-pipeline available at https://github.com/frattalab/rna_seq_single_steps. Custom scripts used to perform differential expression and pathway analysis are available at https://github.com/frattalab/tdp43-apa.

For cross-referencing with differential RNA expression, we used differential expression analysis from cortical-like i3Neurons performed as previously described^13^. Cryptic last exon-containing genes were highlighted if they passed the statistical significance threshold in the Ribo-seq differential expression analysis.

### Analysis of ELK1 transcription factor activity

ELK1 target genes in HeLa cells were accessed from the ChIP-Atlas^83^ on the 15th November 2023. We used the ‘Target genes’ module to obtain a list of target genes that have a ChIP-seq peak within +/− 1 kilobase of transcription start sites. The resulting list contained two HeLa datasets (GSM608163, GSM935326), and was filtered to target genes identified in both datasets Given a reported redundancy of function between ELK1 and other members of the ternary complex factor family^84^ (ELK3 and particularly ELK4), we also attempted to define a unique set of ELK1 target genes. ELK4 target genes in HeLA cells were accessed from ChIP-Atlas on the 29th November 2023 using the same parameters. The resulting list contained 3 HeLa datasets (GSM608161, GSM608162, GSM935351), and we again filtered for target genes identified in all datasets. ELK3 HeLa ChIP-seq data was not available through ChIP-Atlas at the time of publication, and was not considered for further redundancy. ELK3 RNA levels are 10x lower than ELK3 and ELK4 in HeLa TDP-43 knockout cells^39^, so we anticipate this is unlikely to affect our conclusions. ELK1 and ELK4 target gene lists were intersected to define common and unique target genes for each transcription factor. Final target gene lists used are reported in Supplementary Table 5.

RNA-sequencing data from HeLa cells with TDP-43 knockout^39^ were accessed from GSE136366. The data were processed and differential expression was performed as described in Brown et al^10^. Genes were ranked by DESeq2’s test statistic (log_2_ transformed fold change divided by the standard error of the fold change) after removing genes with differential splicing upon TDP-43 knockout, where we can expect to attribute any changes in gene expression to TDP-43 loss of function. Differentially spliced genes were defined using MAJIQ^85^, considering any genes with a probability of > 0.95 as differentially spliced. The target gene sets described above were used as input pathways to fgsea^32^ version 1.24.0 using default settings.

### Code availability

All visualisation and statistical testing was performed in R^86^ version 4.3.2 using the ggplot2^87^ v3.4.4, ggpubr^88^ v0.6.0, ggprism^89^ v1.0.4 and ggrepel^90^ v0.9.4 packages. Preprocessing for visualisation and generation of supplementary tables was performed using tidyverse^91^ v2.0.0, writexl^92^ 1.4.2, and data.table^93^ v1.14. Unless otherwise stated, analyses requiring genomic interval operations or queries with bioinformatics data formats were performed in Python 3.10.11 using PyRanges^73^ 0.0.127, pandas^94^ v2.0.2 numpy^95^ v1.23. Analysis and visualisation code, along with conda^96^ and renv^97^ environments for dependency management, can be accessed at https://github.com/frattalab/tdp43-apa. Alternative repositories for specific analyses are reported in the relevant sections of the methods.

## Author contributions

Conceptualization: PF, MS, SB-S

Data Curation: SB-S, ALB, AM, SB, MZ, OGW, YW, JH

Formal analysis: SB-S, A-LB, PRM, FM, AM, SB

Funding acquisition: PF,MS, MEW, TR

Investigation: SB-S,A-LB,PRM,FM,AM,MY,SB,YA.Q,SH,JNV,KS,ER, MEW

Methodology: SB-S,A-LB,PRM,FM,SE.H,MZ,MK,OGW,JH,NB,MS,PF

Project administration: PF, MS

Resources: JH,MW,PF, TR

Software: SB-S,A-LB,AM

Supervision: PF,MS

Visualisation: SB-S, A-LB, PRM, AM, SB

Writing - Original Draft: SB-S,PF,MS

Writing - Review & Editing: All authors

## Acknowledgements

SB-S is supported by a UK Motor Neurone Disease Association and Masonic Charitable Foundation PhD Studentship (893792). PF is supported by a UK Medical Research Council Senior Clinical Fellowship and the MNDA Lady Edith Wolfson Fellowship (MR/M008606/1 and MR/S006508/1), NIH U54NS123743, Target ALS and The Robert Packard Center for ALS Research. MS is supported by a UKRI Future Leaders Fellowship (MR/T042184/1). PRM is supported by a Wellcome Trust Clinical Training Fellowship (102186/B/13/Z). This work was supported, in part, by the Intramural Research Program of the National Institutes of Neurological Disorders and Stroke (MEW), by the Center for Alzheimer’s and Related Dementias, National Institute on Aging and National Institute of Neurological Disorders and Stroke (MEW), by the Robert Packard Center for ALS Research (MEW).

---

**Supplementary Figure 1-.**
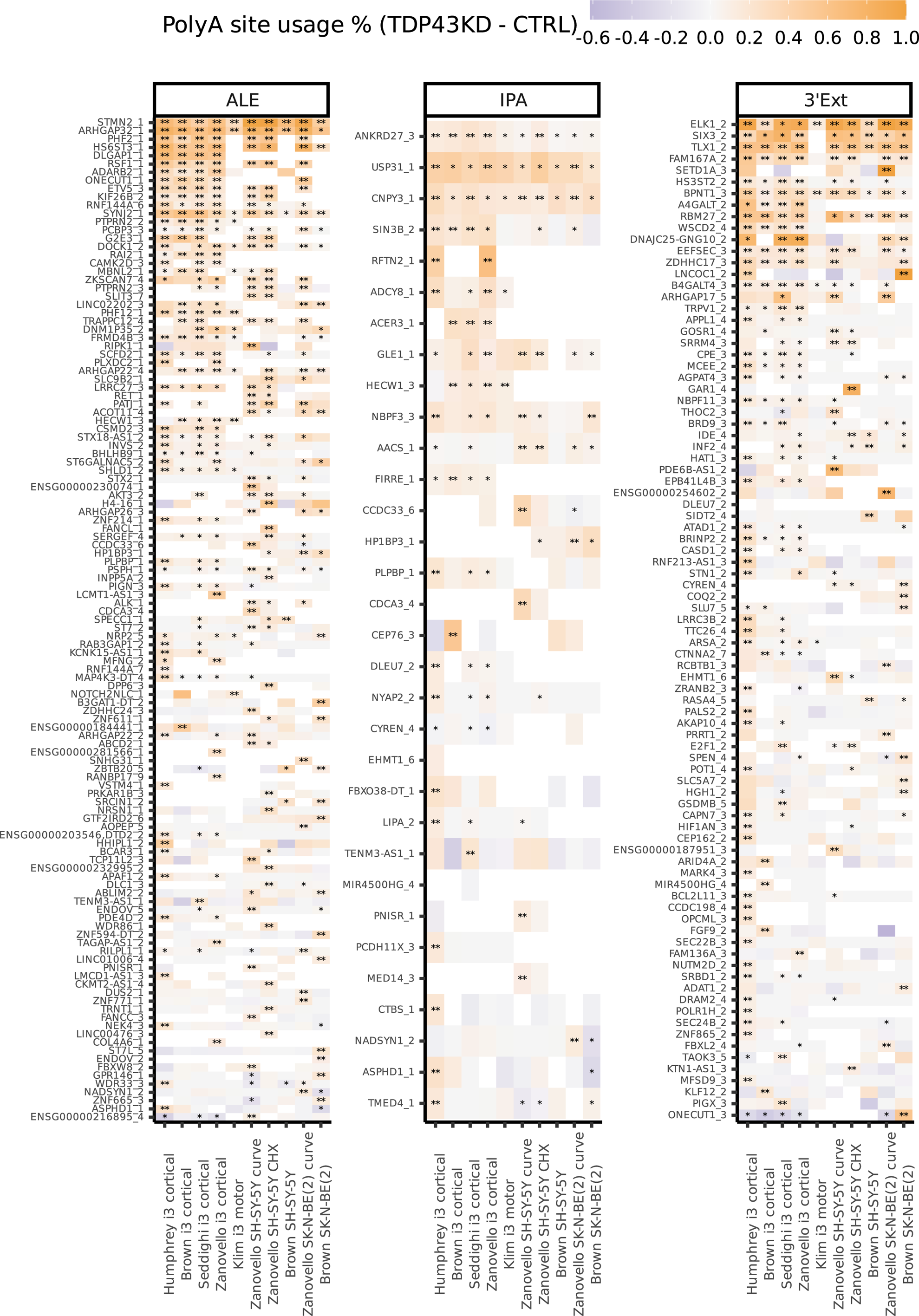
Consistency of response to TDP-43 depletion across compendium of in-vitro datasets Differential usage of cryptic APA events across the compendium of in-vitro datasets. Cells are coloured in accordance to magnitude and direction of change in usage, where positive values (orange) indicate increased usage in TDP-43 knockdown (‘TDP43KD’) samples. Blank cells indicate the event was not expressed at sufficient levels to be assessed for differential usage. Rows are sorted in decreasing order of the sum of −log_10_ transformed p-values weighted by the change in usage between TDP-43 knockdown and control samples (TDP43KD - CTRL) in each dataset. A single asterisk indicates that the isoform was considered significantly regulated in a dataset (Benjamini-Hochberg adjusted p-value < 0.05), and two asterisks indicate the isoform is considered cryptic in a given dataset.

**Supplementary Figure 2-.**
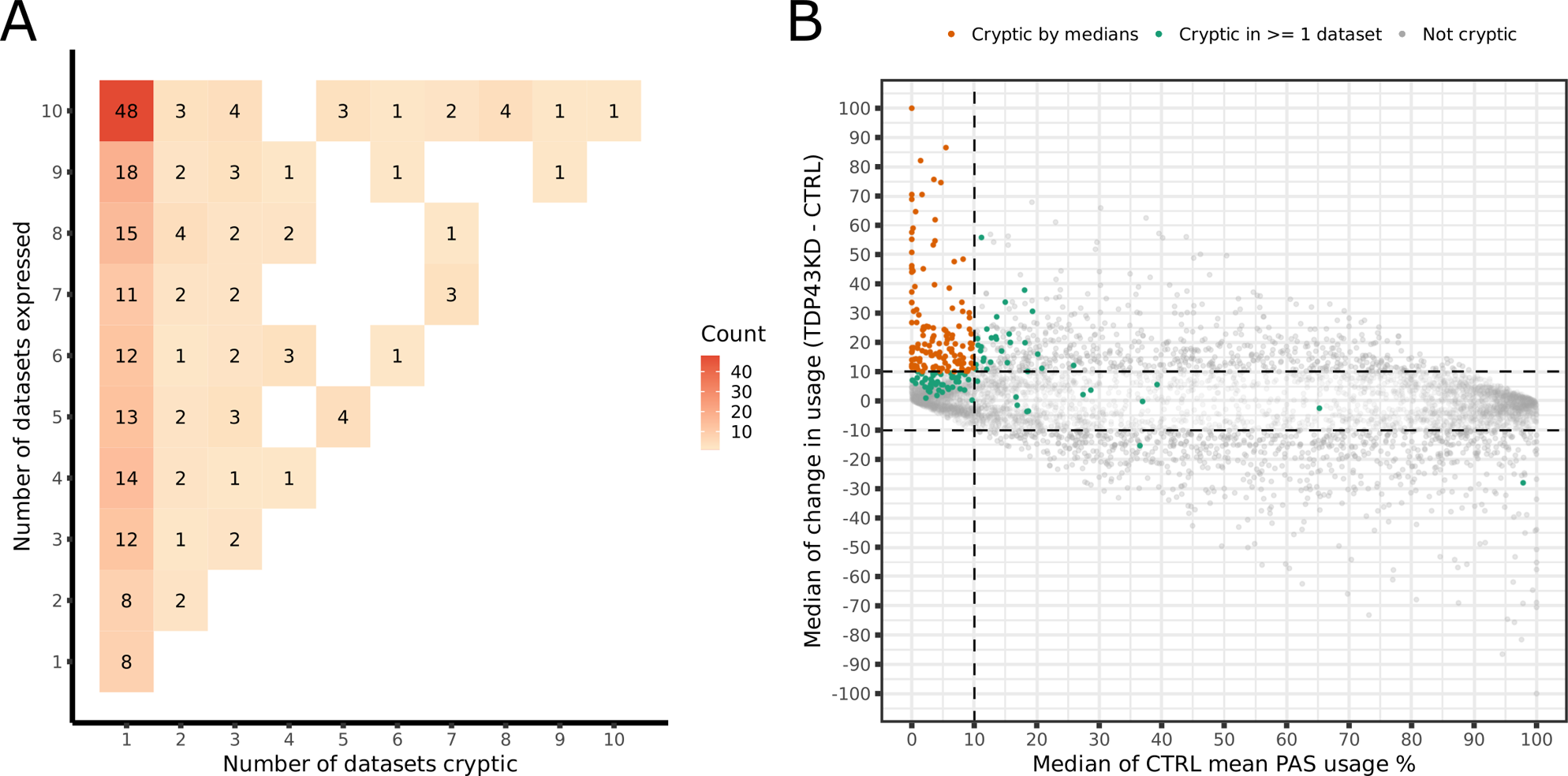
Consistency of cryptic status across compendium of *in vitro* datasets. A) Relationship between the number of datasets in which APAs are called cryptic and their detection. Count labels indicate the number of unique cryptic APAs that fall into the given bin. Events are considered expressed if they pass minimum expression criteria to be evaluated for differential isoform usage (Methods). B) Last exons responsive to TDP-43 depletion. All points represent a last exon passing a Benjamini-Hochberg adjusted p-value < 0.05 threshold in at least one dataset. Where a last exon passes the threshold in multiple datasets, the median values across datasets are calculated to represent the basal usage and change in usage upon TDP-43 depletion. Points that pass cryptic expression criteria in at least one dataset but pass (orange) or fail (green) the criteria when calculating the median change in usage and expression in control (CTRL) cells across datasets with an Benjamini-Hochberg adjusted p-value < 0.05 are highlighted.

**Supplementary Figure 3-.**
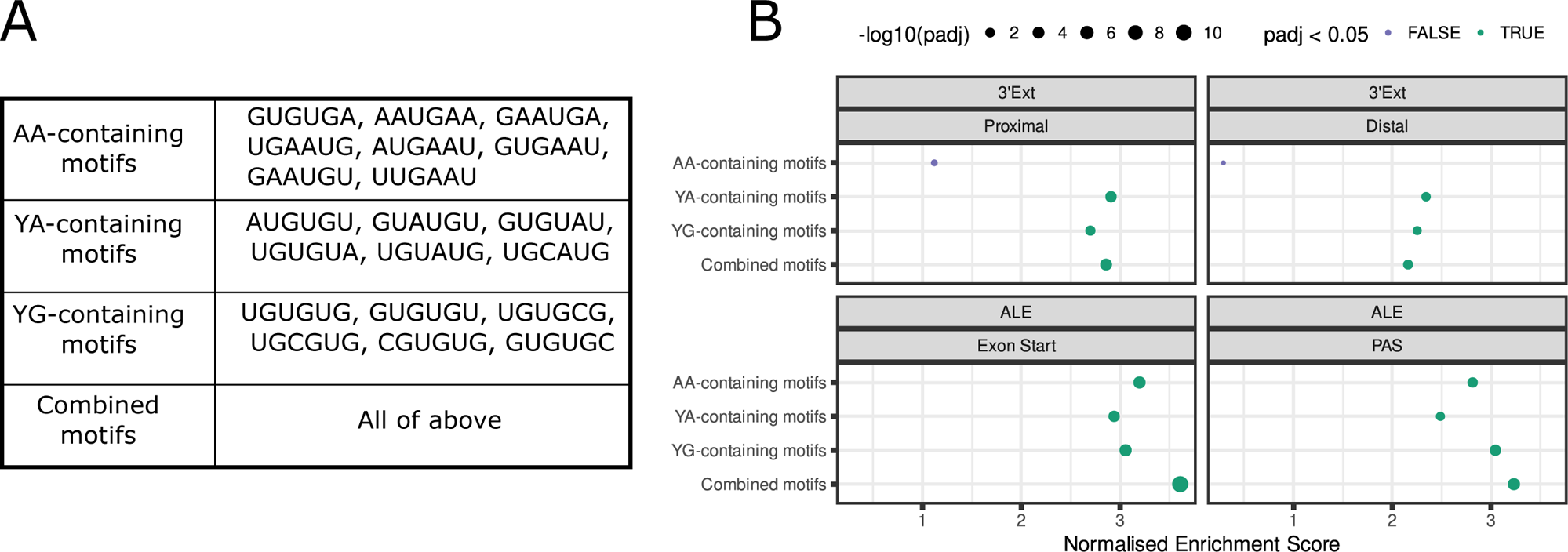
Enrichment of previously defined TDP-43 binding hexamers at cryptic APA boundaries. A) Table listing previously defined TDP-43 hexamer groups^6^. ‘Y’ codes for a pyrimidine nucleotide. B) Gene set enrichment analysis (GSEA) of enriched TDP-43 binding 6mers on de-novo enriched 6-mers around cryptic landmarks. The panels and labels correspond to regions evaluated for iCLIP binding as in Fig. 1D. The area of the points is proportional to the −log_10_ transformed adjusted p-value (adjusted with respect to all region types and motif groups), and the colour denotes whether the Benjamini-Hochberg adjusted p-value passes (green) or fails (purple) a significance threshold of < 0.05.

**Supplementary Figure 4-.**
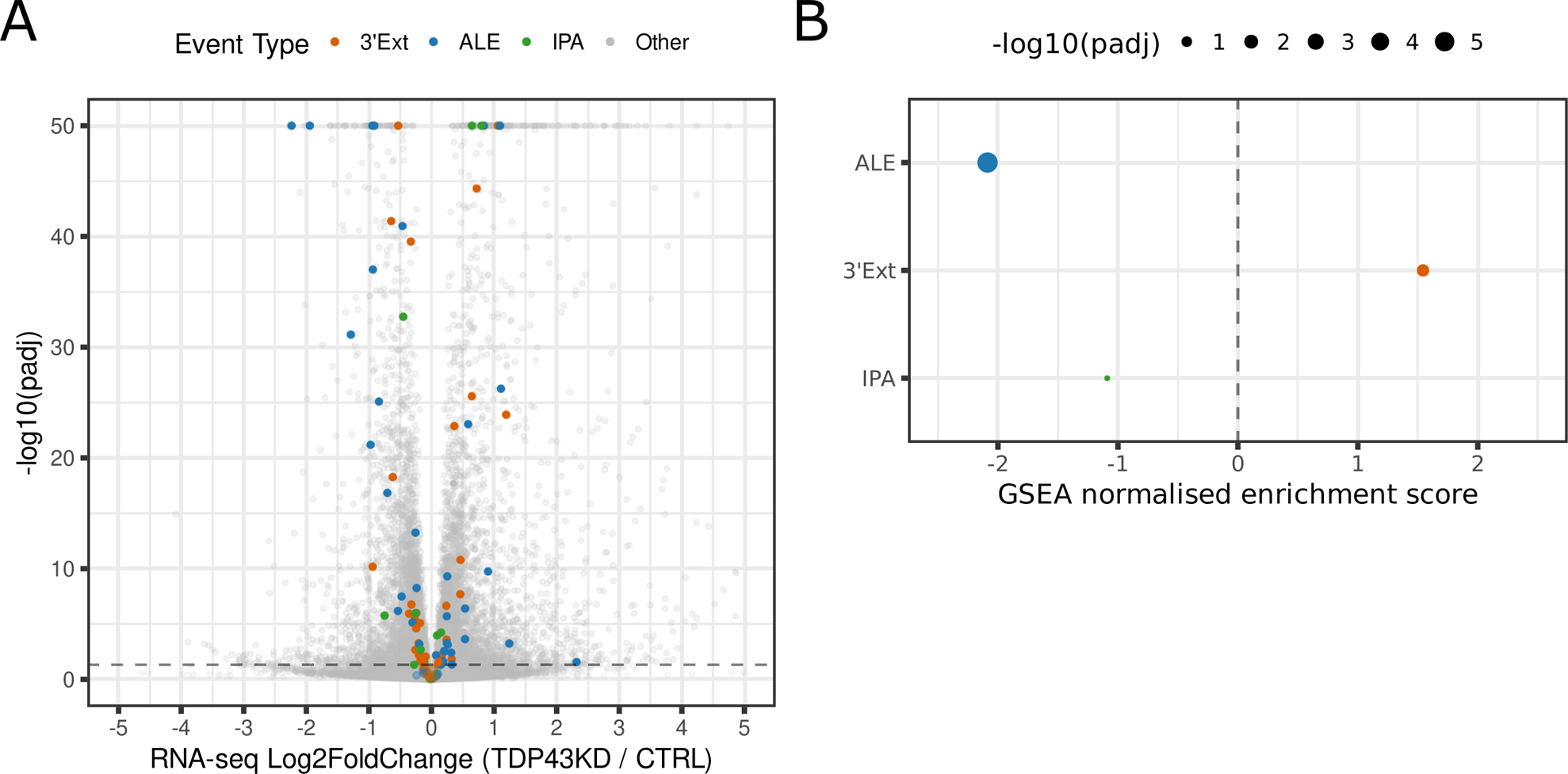
Analysis of cryptic APA categories in Ribo-seq data. A) Volcano plot of differential expression analysis of RNA-seq data between TDP-43 knockdown (TDP43KD) and control (CTRL) i3Neurons. Cryptic APA genes with significant differential expression (Benjamini-Hochberg adjusted p-value < 0.05) are highlighted in orange (3’Ext), blue (ALE) or green (IPA). Genes with a −log_10_ transformed Benjamini-Hochberg adjusted p-value greater than 50 are collapsed to 50 for visualisation purposes. B) Gene Set Enrichment Analysis (GSEA) of cryptic APA categories in i3Neuron Ribo-seq differential expression fold change ranks. The area of the points is proportional to the −log_10_ transformed Benjamini-Hochberg adjusted p-value. Points are coloured according to their APA category as in A).

